# Compensation Strategies for Bioelectric Signal Changes in Chronic Selective Nerve Cuff Recordings: A Simulation Study

**DOI:** 10.1101/2020.11.12.380121

**Authors:** Stephen Sammut, Ryan G.L. Koh, José Zariffa

**Affiliations:** Institute of Biomedical Engineering, University of Toronto, Toronto, ON M5S 3G4, Canada; KITE, Toronto Rehab, University Health Network, Toronto, ON, M5G 2A2, Canada; Edward S Rogers Sr Department of Electrical and Computer Engineering, University of Toronto, Toronto, ON M4G 3V9, Canada; Rehabilitation Sciences Institute, University of Toronto, Toronto, ON M5S 2E4, Canada

## Abstract

Peripheral nerve interfaces (PNIs) allow us to extract motor, sensory and autonomic information from the nervous system and use it as control signals in neuroprosthetic and neuromodulation applications. Recent efforts have aimed to improve the recording selectivity of PNIs, including by using spatiotemporal patterns from multi-contact nerve cuff electrodes as input to a convolutional neural network (CNN). Before such a methodology can be translated to humans, its performance in chronic implantation scenarios must be evaluated. In this simulation study, approaches were evaluated for maintaining selective recording performance in the presence of two chronic implantation challenges: the growth of encapsulation tissue and rotation of the nerve cuff electrode. Performance over time was examined in three conditions: training the CNN at baseline only, supervised re-training with explicitly labeled data at periodic intervals, and a semi-supervised self-learning approach. This study demonstrated that a selective recording algorithm trained at baseline will likely fail over time due to changes in signal characteristics resulting from the chronic challenges. Results further showed that periodically recalibrating the selective recording algorithm can maintain its performance over time, and that a self-learning approach has the potential to reduce the frequency of recalibration.

## 1. Introduction

The peripheral nervous system controls the flow of motor, sensory, and autonomic information from the central nervous system to other parts of the body and therefore exerts control over a multitude of physiological functions. Bioelectric signals recorded from the nervous system can be used to create commands for neuroprosthetic devices (which interact with the nervous system to restore lost function) and to regulate the activity of the nervous system through closed-loop neuromodulation. Examples of peripheral nerve recordings in neuroprosthetic applications include decoding motor signals to control a computer cursor [1], [2] or prosthetic limb [3] [4], or using sensory feedback to control a functional electrical stimulation system [5], [6]. Examples in neuromodulation applications include regulating blood pressure through selective vagus nerve stimulation [7], as well as using the electroneurogram of the vagus nerve to detect the onset of a seizure [8], [9]. Currently, a major roadblock to closed-loop neuroprosthetic and neuromodulation applications is the need for techniques that can extract useful information from low-amplitude, noisy recordings [10].

Of the available peripheral nerve electrode designs [11], nerve cuff electrodes are an appealing choice as they have a relatively long history of use [12]–[14] and have been shown to be safe for long-term or chronic implantation both in animals [15]–[21] and humans [6], [22]. However, these devices are extraneural (i.e. do not penetrate the nerve), resulting in low signal-to-noise ratio (SNR) and recording selectivity.

As such, a variety strategies have been developed in order to improve upon the selectivity and performance of these interfaces, for example, cuff electrodes with multiple sites [23]–[26], intricate stimulation modes [23], [27], and implementation of various signal processing algorithms [28], [29]. Several research groups have attempted to improve the recording selectivity of cuff electrodes by using multi-contact nerve cuff configurations. For example, conduction velocity information has been determined by investigating the propagation of the neural signal across multiple contacts along the length of the nerve [30]–[34]. Alternatively, a number of studies have attempted to achieve spatial selectivity of neural recordings using beamforming [15], [35]–[37] or source localization methods [38], [39]. Building on the results above, Koh et al. used spatiotemporal signatures to characterize the recordings from a nerve cuff with an array of contacts [40]–[42]. By using these spatiotemporal signatures as input to a convolutional neural network (CNN), individual natural compound action potentials (nCAPs) could be classified according to the neural pathway that produced them [42]. This CNN is referred to as the extraneural spatiotemporal compound action potential extraction network, or ESCAPE-NET. In this manner, the firing patterns of different pathways could be reconstructed, thus improving recording selectivity compared to previous approaches.

Although ESCAPE-NET has achieved promising results in acute experiments, its response to the functional and morphological changes that occur during chronic implantation must be investigated before it can be translated to human subjects. Here we used a simulation study to characterize the changes in the performance of the selective recording algorithm over time, and to evaluate strategies to compensate for these changes. In this study, two major factors likely to cause bioelectric signal changes in recordings of multi-contact nerve cuff electrodes, and therefore reduce selectivity, were simulated: 1) the buildup of encapsulation tissue surrounding the nerve, which is an immune response to an implanted neural electrode and causes changes in the electrical interface [43]–[45] and 2) shifts in the position of the nerve cuff electrode as a result of physical movement, known to impact recording selectivity [46], [47]. After simulating and characterizing the changes that occurred over time, we compared three approaches for updating ESCAPE-NET during chronic implantation in order to maintain high levels of performance.

## 2. Methods

The first step in our study was to simulate recordings from a multi-contact nerve cuff electrode implanted on a rat sciatic nerve. We then altered the model to simulate signal changes during chronic implantations. Lastly, methods to compensate for those changes over time were compared.

### 2.1. Model construction

A previously described finite element model of the rat sciatic nerve and nerve cuff electrode in a saline bath [46] was used as the basis for our simulations in this study. The model was originally developed using MRI imaging of the nerve and includes progressive branching of the sciatic nerve into the tibial, peroneal, and sural nerves. The anatomy of this model was manually segmented using Seg3D [48], before implementing the modifications described below.

The endoneurium and epineurium were manually traced and thus accurately reflect the physical dimensions of these structures, while the perineurium was added around each fascicle with an approximate thickness of 0.065 mm, due to the fact that thinner layers have previously been found to cause errors in the meshing process [46].

Once segmentation of the rat sciatic nerve was completed and verified, it was exported to MATLAB, where a tetrahedral mesh model was generated using Iso2Mesh [49]. This baseline is shown in Figure 1. The layout of the nerve cuff electrode is shown in Figure 2.

**Figure 1:**
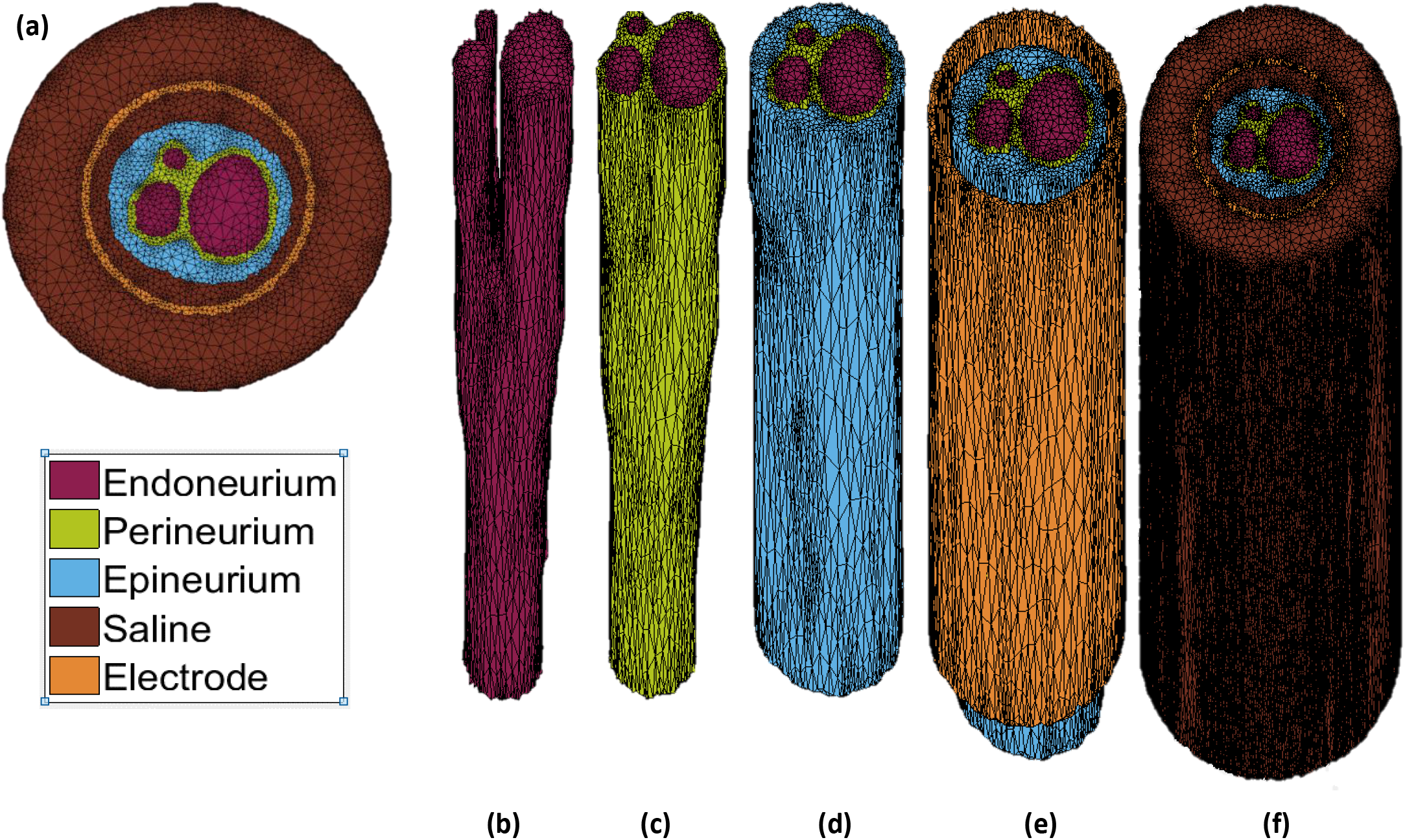
Finite Element (FE) tetrahedral mesh model. (a) Top-down view of the base FE mesh model. Individual layers of the FE mesh model (b) endoneurium, (c) perineurium, (d) epineurium, (e) cuff electrode, and (f) saline.

**Figure 2:**
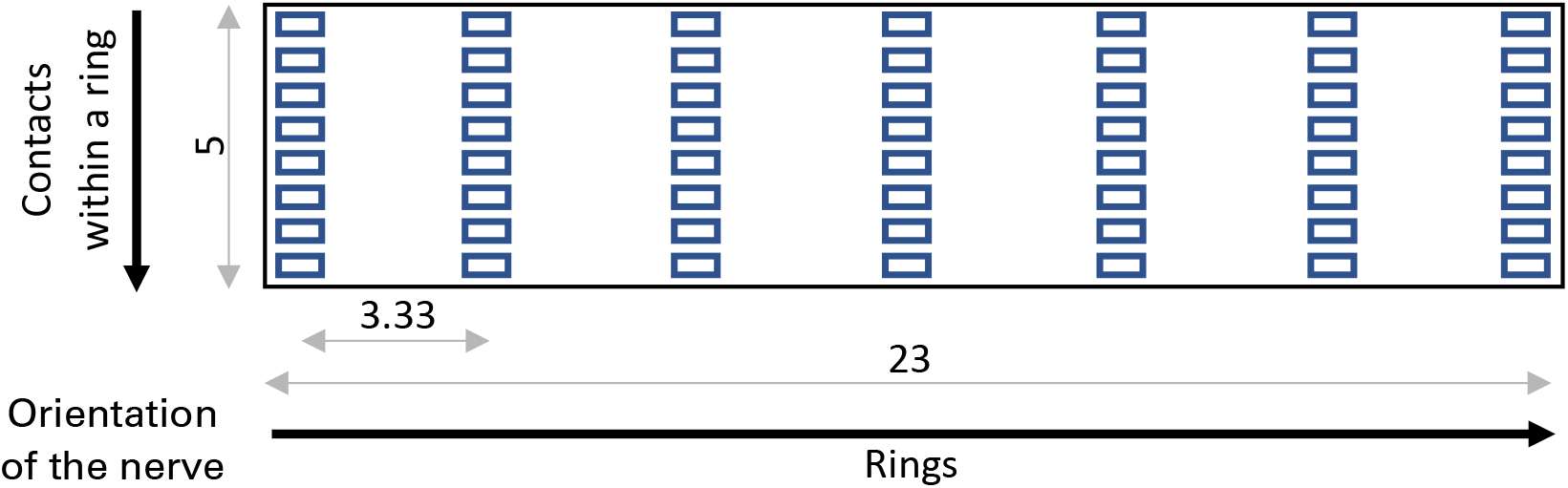
Visualization of the unrolled nerve cuff electrode design used in modelling, with dimensions shown (millimeters).

Following the generation of the mesh model, finite element analysis was conducted in SCIRun [50] in order to generate a leadfield matrix. The leadfield, represented by a matrix **L** with entries (*i, j*), characterizes the influence that a unit current dipole positioned at a mesh element *j* has on the electrical potential observed at an electrode contact *i*. **L** is of size M x N, where M represents the number of electrode contacts and N represents the number of elements which form the tetrahedral mesh. The leadfield matrix was constructed according to the process described by Weinstein et al. [51].

Parameters of the tetrahedral mesh model, including conductivity values and structure dimensions, are summarized in Table 1, and are established based on similar models described in the literature.

**Table 1:**
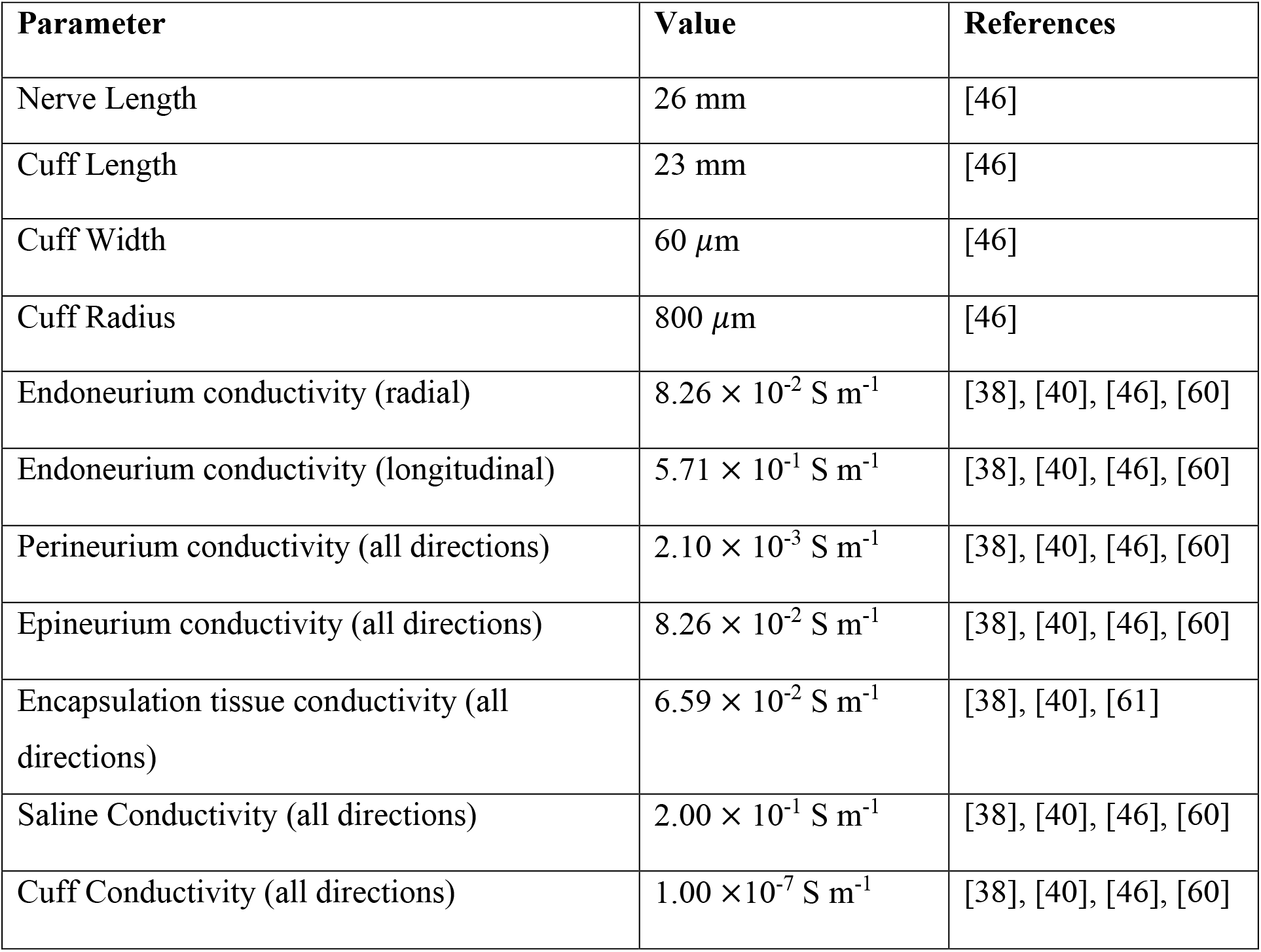
FE mesh model parameters.

| Parameter | Value | References |
| --- | --- | --- |
| Nerve Length | 26 mm | [46] |
| Cuff Length | 23 mm | [46] |
| Cuff Width | 60 $\mu\text{m}$ | [46] |
| Cuff Radius | 800 $\mu\text{m}$ | [46] |
| Endoneurium conductivity (radial) | $8.26 \times 10^{-2} \text{ S m}^{-1}$ | [38], [40], [46], [60] |
| Endoneurium conductivity (longitudinal) | $5.71 \times 10^{-1} \text{ S m}^{-1}$ | [38], [40], [46], [60] |
| Perineurium conductivity (all directions) | $2.10 \times 10^{-3} \text{ S m}^{-1}$ | [38], [40], [46], [60] |
| Epineurium conductivity (all directions) | $8.26 \times 10^{-2} \text{ S m}^{-1}$ | [38], [40], [46], [60] |
| Encapsulation tissue conductivity (all directions) | $6.59 \times 10^{-2} \text{ S m}^{-1}$ | [38], [40], [61] |
| Saline Conductivity (all directions) | $2.00 \times 10^{-1} \text{ S m}^{-1}$ | [38], [40], [46], [60] |
| Cuff Conductivity (all directions) | $1.00 \times 10^{-7} \text{ S m}^{-1}$ | [38], [40], [46], [60] |

### 2.2. Simulation of Chronic Factors

#### 2.2.1. Growth of Encapsulation Tissue

The original model was modified in order to incorporate encapsulating tissue into the segmentation as an additional material type. This new structure was incorporated into the original segmentation between the epineurium and cuff electrode throughout each layer, as illustrated in Figure 3. This process was repeated multiple times in order to generate a total of four models with varying degrees of encapsulation tissue growth(referred to as ET1, ET2, etc.), and therefore simulate a number of time points following the implantation of the nerve cuff electrode. In this simulation, time elapsing was simulated through continuous growth of encapsulation tissue between models. Note that in reality, encapsulation tissue will eventually also surround the cuff. However, the external layer of tissue is unlikely to affect our simulated measurements at the recording contacts, which are located on the inner surface of the cuff, and therefore was neglected in our modeling.

**Figure 3:**
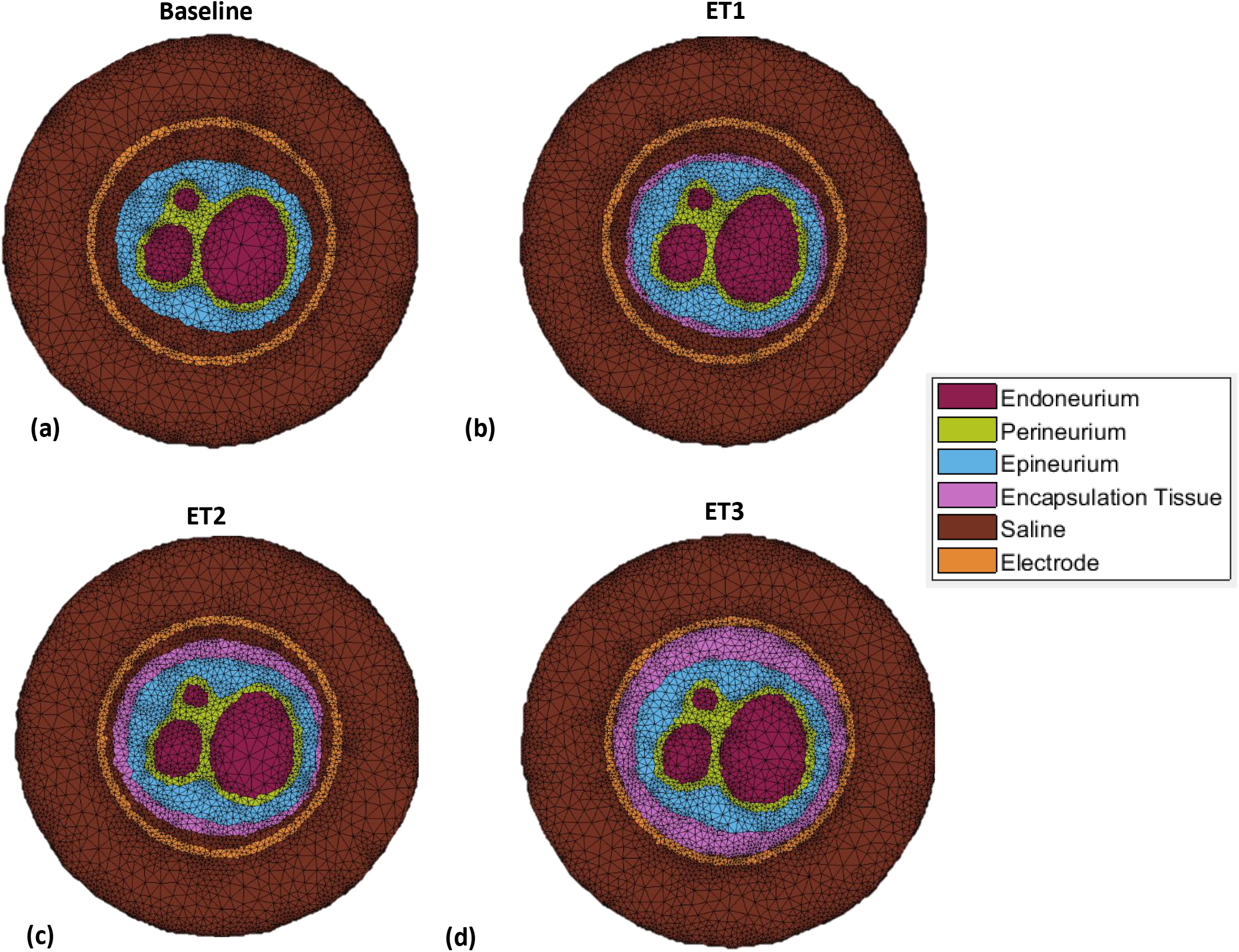
Top-down view of FE mesh models from: (a) no encapsulation tissue to (d) full encapsulation of the nerve.

#### 2.2.2. Rotation of the Nerve Cuff Electrode

In order to simulate the rotation of the nerve cuff electrode around the rat sciatic nerve, the electrode positions were rotated in three-dimensional space around the central axis of the nerve. Given the geometry of this nerve cuff electrode, eight contacts in each ring surround the perimeter of the nerve. As such, a rotation of 45° (360°/8 contacts = 45°) from the original position results in all electrode contacts being placed in the starting position of a different contact. For this reason, the 45° was selected as the maximal rotation, and increments of 5° were simulated. A new leadfield was generated for each rotation, using the new contact locations. In this simulation, deviation from the original signals was simulated through continuous rotation of the nerve cuff electrode.

### 2.3. Simulated Recordings

Myelinated mammalian fiber action potentials were obtained with the *Neurocal* MATLAB toolbox (Case Western University, Ohio, USA) using the Schwarz–Reid–Bostock model [52]. This model is typically used to simulate single fiber action potentials, however, a simplifying assumption was made here that it could be used to approximate the shape of a CAP; this assumption corresponds to the simplified CAP scenario of multiple nerve fibers firing in perfect synchrony, leading to a version of the single fiber action potential with scaled-up amplitude, as used in previous modeling [40].

The CAPs were propagated through the model at differing spatial locations within each of three neural pathways, by selecting mesh elements from within a given fascicle. Multiple points for a single pathway were manually defined throughout the model starting from the proximal end and moving towards the distal end. MATLAB was used in order to generate a spline interpolation along the entire length of the nerve from the provided points, thus defining a neural pathway [46]. This procedure was performed for the tibial, peroneal, and sural fascicles.

The positioning of the Nodes of Ranvier along a neural pathway was defined according to the fiber type being simulated. In this simulation, the spacing of the Nodes Ranvier were 1.70 mm for Aα fibers (tibial and peroneal pathways) and 1.16 mm for Aβ fibers (sural pathway).

Conduction velocities of 94.86 m/s and 64.72 m/s were used for the Aα and Aβ fibers, respectively [40]. These parameters were used in conjunction with the leadfield matrix to simulate all 56-channel measurements of a CAP propagating through a single neural pathway. The simulated recordings were then tripole referenced using the average of all outer ring contacts [53].

By combining CAPs from a single neural pathway, a time series was created. A single time series was created for each of the three fascicles (tibial, peroneal, and sural), composed of 10,000 CAPs within a duration of 53.333 seconds. These specifications were chosen in order to add noise to the simulated recordings in a consistent manner to Koh et al. [40]; hereafter, CAPs are analyzed individually without consideration of firing rates. White Gaussian noise was then added to the time series in order to simulate baseline and instrumentation noise commonly observed in these types of recordings *in vivo*, using decreasing levels of signal-to-noise ratios (−5, −10, and − 15 dB), defined using the measured signal power over the entire time series.

### 2.4. CAP Classification

#### 2.4.1. Datasets

The simulated CAPs were individually classified according to the neural pathway used to generate the recording (tibial, peroneal, or sural fascicle). Therefore, classification here was a three-class problem.

Once the noisy time series were created, individual CAP spatiotemporal signatures were extracted, based on peak detection prior to the addition of noise. Signatures 100 time samples long (3.333 ms, using a sampling rate of 30 kHz) were created by obtaining 49 points before and 50 points after the location of the CAP peak, for each of the 56 channels. This data was then normalized within the range of −1 to 1 based on the maximum amplitude seen across all CAPs. These data points served as either training or testing samples, depending on the strategy being evaluated, and were provided to our CNN (ESCAPE-NET) with their respective labels based on the neural pathway producing them. A dataset containing a total of 30,000 CAPs (10,000 CAPs x 3 pathways) was created for each model in each of the two simulations (encapsulation tissue growth and rotation of the cuff electrode).

#### 2.4.2. Convolutional Neural Network Convolutional Neural Network

##### Convolutional Neural Network Convolutional Neural Network

A previously described CNN, ESCAPE-NET [42], was used to classify the spatiotemporal signatures produced by the CAPs in the multi-contact nerve cuff. As such, the inputs to ESCAPE-NET were 56×100 spatiotemporal signatures created as described above. The CNN used in this work was constructed with Python 2.7 using the Keras application programming interface (API) with TensorFlow [54] as the backend.

The spatiotemporal signature used as input to ESCAPE-NET can exhibit different patterns depending on the ordering of the contacts. Alternatively, the network can be provided with more than one such representation, in order to exploit possibly complimentary information [42]. In the case of this project, ESCAPE-NET’s dual input version was used. In this version of the network, the architecture is split into two identical sides in order to process two input configurations obtained from the matrix of the multi-contact nerve cuff electrode recordings (contacts ordered by rings or longitudinally, referred to as “spatial emphasis” or “temporal emphasis”, respectively). A block diagram of ESCAPE-NET’s architecture can be seen in Figure 4, while additional details can be found in [42].

**Figure 4:**
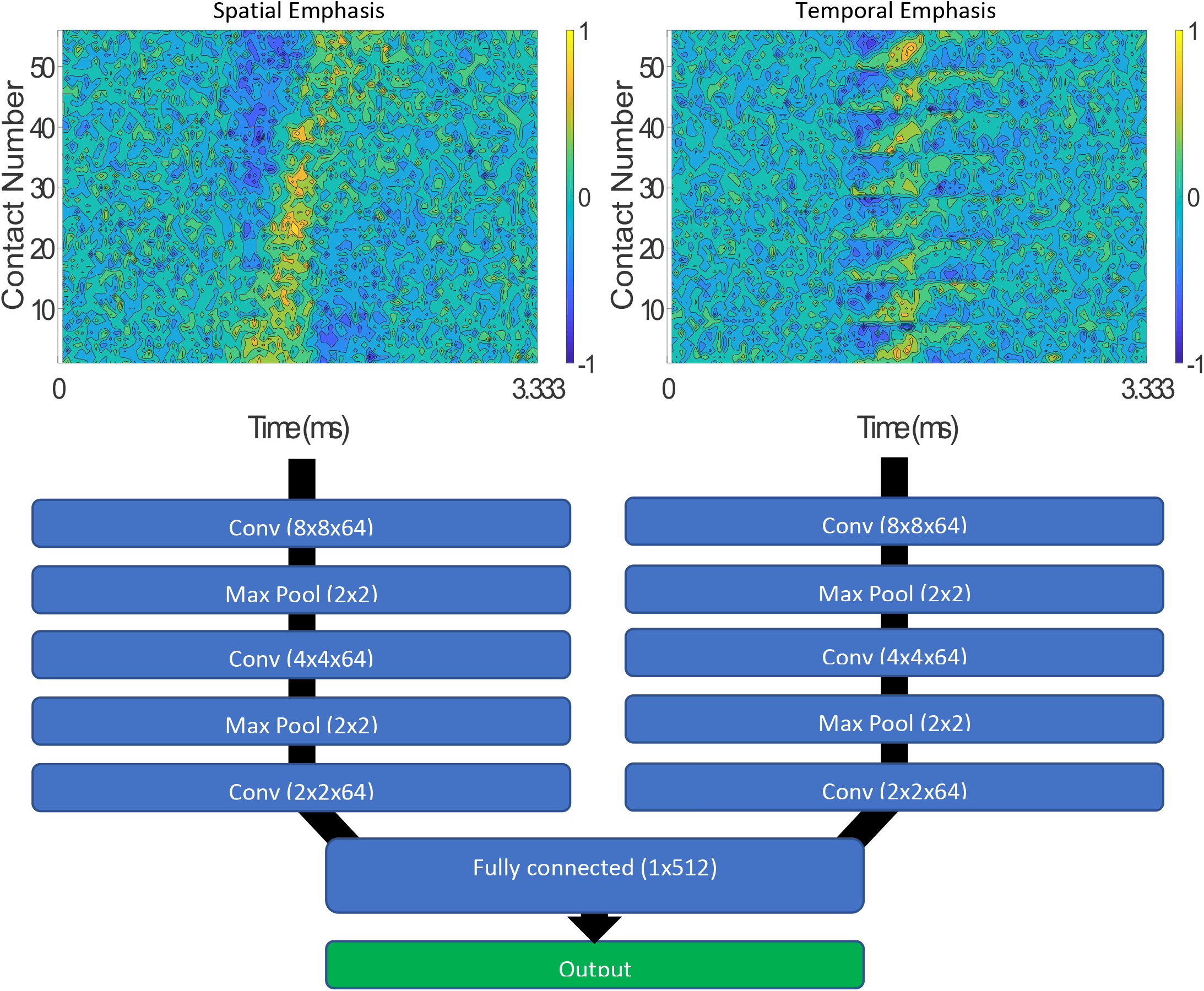
ESCAPE-NET architecture with Spatial and Temporal Emphasis representations of the multi-contact nerve cuff recordings (SNR = −5 dB) shown as input to the network [42].

### 2.5. Classifier Update Strategies

When applying the CNN over the multiple simulated time points, one of three training approaches was used: 1) Baseline Calibration (control condition), 2) Periodic Recalibration, and 3) a Self-Learning approach.

Evaluation was achieved in all three of the approaches using five-fold cross validation. In order to create a training and testing set for the convolutional neural network, the 30,000 CAPs generated for each model were split into five folds, with each fold containing 6,000 CAPs (2,000 CAPs from each of the three pathways). Each fold was used in turn as the testing set for cross-validation, in order to evaluate each of the strategies at each time point. The analysis was repeated separately for each of the 3 noise levels. The classification performance of each network at each time point was assessed by calculating the macro F_1_-score (i.e. calculating the F_1_-score for each class and averaging them).

#### 2.5.1. Baseline Calibration

In the first approach, the classifier was trained once, at baseline, representing the initial implantation of the nerve cuff electrode with no encapsulation tissue or rotation present. The CNN was evaluated on data from subsequent time points without further modification. In this way, we quantified the performance of the classifier over time as factors likely to cause bioelectrical signal drift begin to arise during chronic implantation. This simulation represents a scenario in which a neural interface is calibrated at the time of initial implantation, and its performance evaluated over time without additional recalibration.

#### 2.5.2. Periodic Recalibration

In the second approach, the network, again initially trained at baseline, was retrained at future points in time with new, explicitly labelled, data. At each new time point, the network was first initialized with the final learned weights of the previous time point, then retrained using an explicitly labelled datasets generated by the model for the current time point. This process was repeated for each subsequent point in time. This simulation represents a scenario in which a user carries out an explicit recalibration procedure at regular intervals.

To evaluate the periodic calibration approach, the supervised network was retrained at each new time point with four folds (24,000 CAPs) of the new time point data, and then tested using the remaining fold (6,000 CAPs). This process was repeated using each of the 5 cross-validation folds in turn as the test set.

#### 2.5.3. Self-Learning Approach

In the final approach, the network, again initially trained at baseline, was retrained at future time points without the need for manual labelling of new data. This approach is based on methods described by Schwemmer et al. [55] and utilizes “predicted” data. At each new time point, new data was first classified using the network obtained in the previous time point. Only data samples predicted by the network with high-enough confidence (95% confidence) were chosen, along with their estimated labels, in order to create what is referred to as a “self-labelled” dataset (see Figure 5). The network was then initialized with the final learned weights of the previous time point and retrained using the self-labelled dataset in order to update its weights. This process was repeated for each subsequent point in time. This simulation represents a scenario in which an implanted neural interface is recalibrated periodically, without the need for user or experimenter intervention.

**Figure 5:**
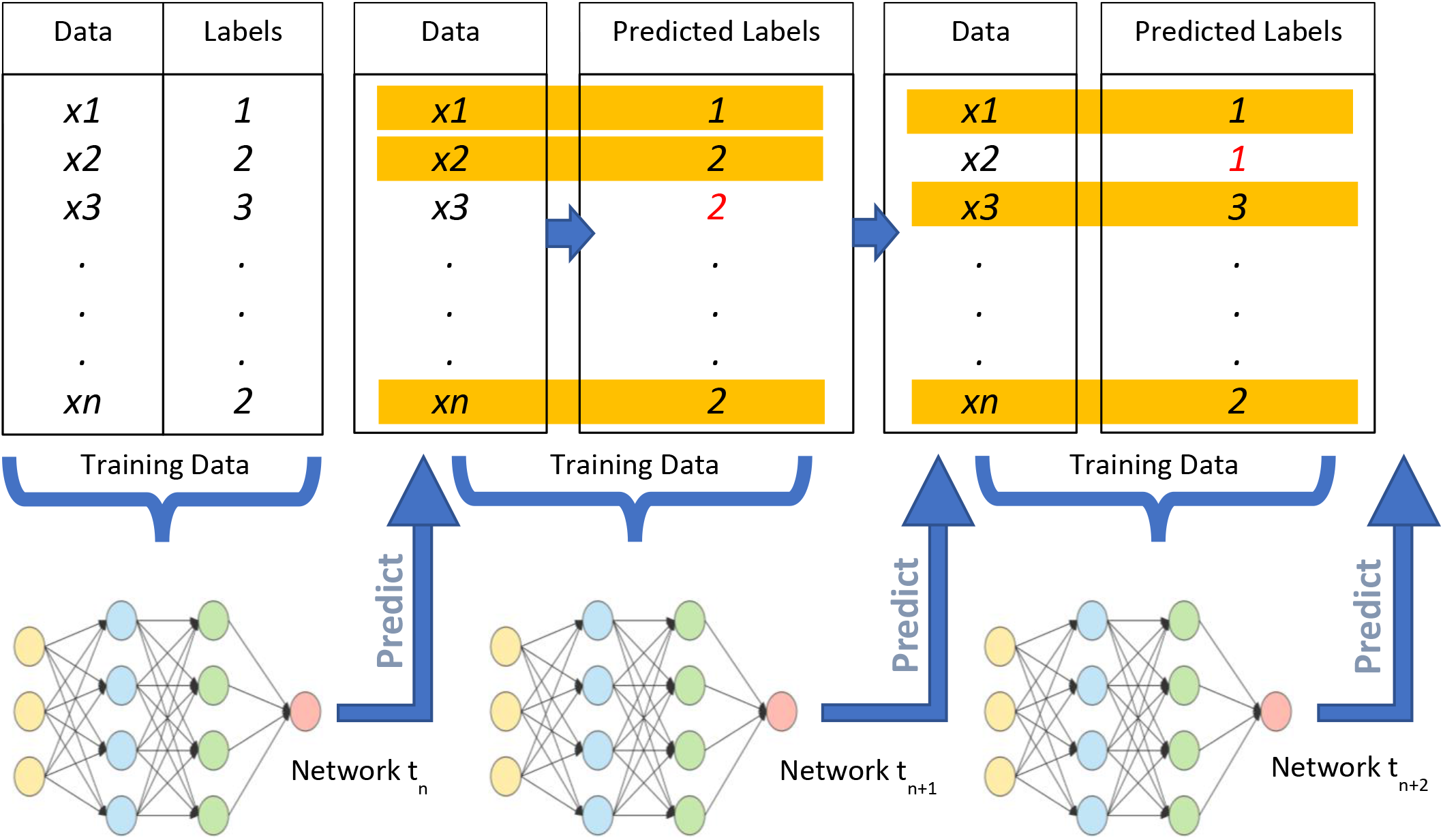
Visual representation of the processes involved in self-learning approach. In each step, the previously obtained model is used to predict labels at the new time point. The network is then updated/retrained using these predicted datasets, at the current point in time. Here, the highlighted text represents correctly predicted samples, while red text represents mislabeled samples, both of which may be incorporated into a “self-labelled” dataset depending on the confidence level.

To evaluate the self-learning approach, the network at each new time point was first initialized using the learned weights from the previous time point. The classifier was retrained with four folds of the new time point data, labelled by the network itself. The network was then tested using the remaining fold, but with the explicit labels provided. The process was repeated five times with different testing folds for cross validation.

The classification of the network achieved using each of the three updating strategies was assessed by calculating the macro F_1_-score (i.e. calculating the F_1_-score of each class and then averaging them).

A timeline of the training and testing of the CNN involved in each of the three update strategies, using the growth of encapsulation tissue simulation as an example, is depicted below in Figure 6.

**Figure 6:**
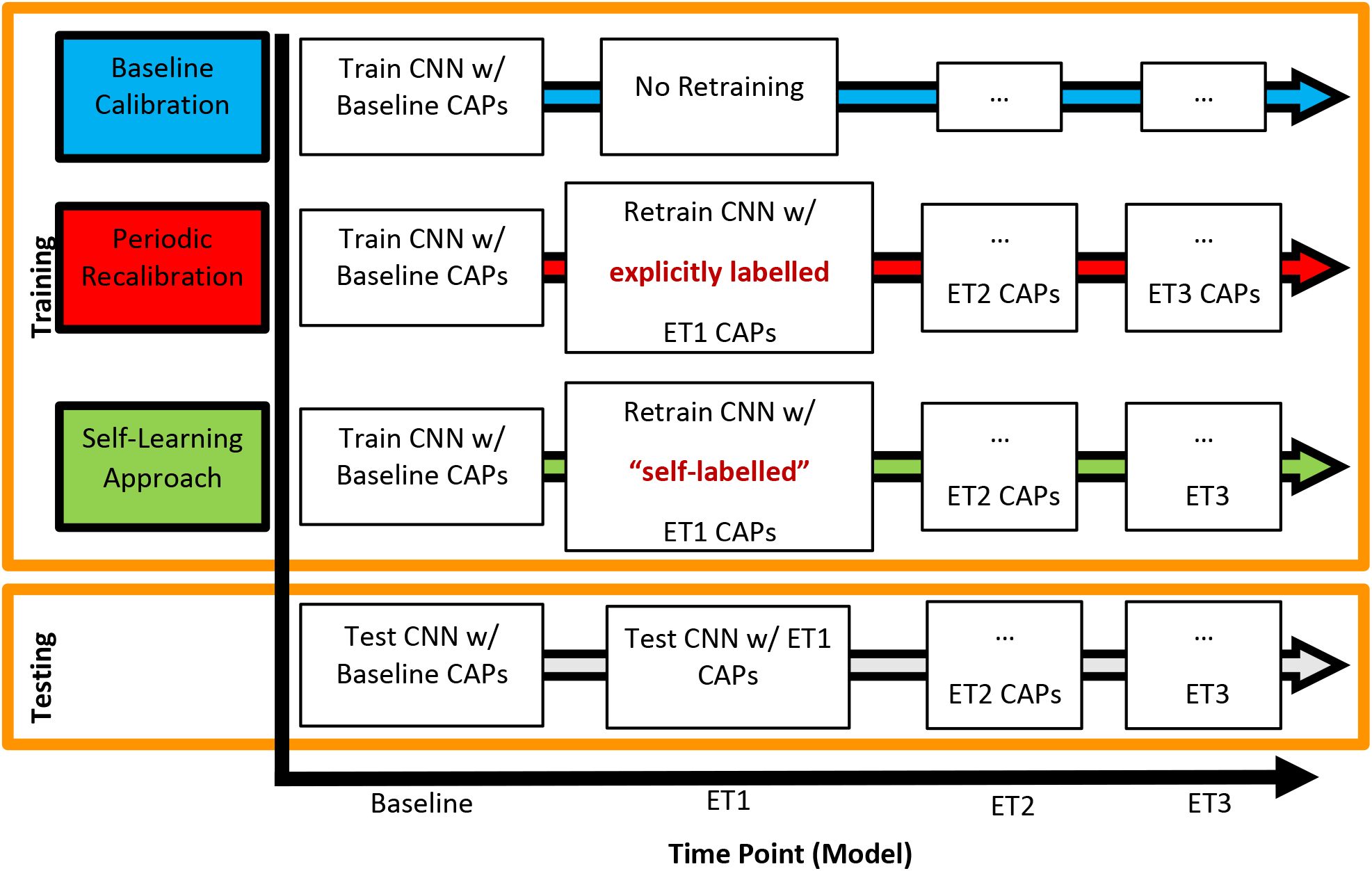
Training and testing scenarios of the three updating approaches. The vertical axis of the figure shows the update strategy, while the horizontal axis represents time elapsing and subsequent changes in the interface. Text boxes within the figure show training and testing approaches corresponding to specific update strategies at a given time point (in this case representing the growth of encapsulation tissue).

#### 2.5.4. Further Analysis of the Self-Learning Approach

Due to the nature of the self-learning approach, several factors are likely to influence the efficacy of this machine learning update strategy. We conducted additional analyses of the encapsulation tissue simulation to characterize the impact of the following parameters.

##### 2.5.4.1. Training frequency

First, we hypothesize that the greater the time between training intervals, the less effective the self-learning method will be. This hypothesis is based on the assumption that a relatively long interval between training times will correspond to greater changes in the recorded signals. Large signal changes will lead to poorer performance of a classifier trained at the previous time point, which in turn may cause inaccurate predicted labels and a degradation in classification accuracy. As such, updating the network at less frequent time points will likely result in poorer performance over time. We therefore investigated the efficacy of the self-learning approach in scenarios where the time between training intervals was modified.

In order to investigate whether the self-learning approach would benefit from more frequent training of the CNN, linear interpolation was used between the CAPs from successive time points in the finite element models in order to generate data from additional time points. This approximation was chosen as it would be quite difficult to create additional three-dimensional mesh models with smaller more gradual increases in encapsulation tissue growth due to space limitations in the segmentation step and the need for finer meshes. In this experiment, the self-learning approach was implemented as described above, and the network was then retrained twice, four-times, and eight-times as often as the other two update strategies.

##### 2.5.4.2. Initial Performance Level

Another factor that could affect the efficacy of the self-learning approach is the initial performance level. If the performance of the network from the previous time point is relatively high, it may more correctly predict and thus label samples in the new time point. Conversely, an inaccurate network will predict incorrect labels, leading to error accumulation and performance degradation. As such, the performance of the network at one time point will affect the number and accuracy of the labelled samples at the next time point. We therefore investigated the efficacy of the self-learning approach in scenarios where the initial performance level was modified.

To this end, we needed a set of different initial performance levels. This was accomplished using the results achieved at ET1 for different updating methods and SNRs. Thus, for this portion of the analysis, ET1 was used as the starting point. Specifically, six different initial starting performances were achieved by initializing the network with weights obtained at baseline at each noise level (−5, −10, and −15 dB) and then applying either the periodic recalibration approach or the self-learning approach at ET1.

##### 2.5.4.3. Evaluation

Different training frequencies (1x, 2x, 4x, 8x) were applied for each starting level at each SNR. The purpose of this analysis was to understand how the self-learning approach performed compared to a no-update control, depending on the initial performance and retraining frequency. The control was a model trained at ET1 for each of the initial performance levels and applied to subsequent time points without retraining.

The primary metric for this analysis was the slope of the F_1_-score between ET1 and ET2, compared to the slope of the control. For example, a method that led to a higher performance than the control at ET2 would have a less negative slope, and thus could be said to be beneficial, i.e. to have slowed the decline in performance over time.

## 3. Results

### 3.1. Simulated Recordings

Once a compound action potential has been propagated through the model, measurements at all 56 channels of the simulated nerve cuff electrode were obtained, as illustrated in Figure 7a (shown with no noise added). Examples of time series segments with different noise levels are shown in Figure 7b.

**Figure 7:**
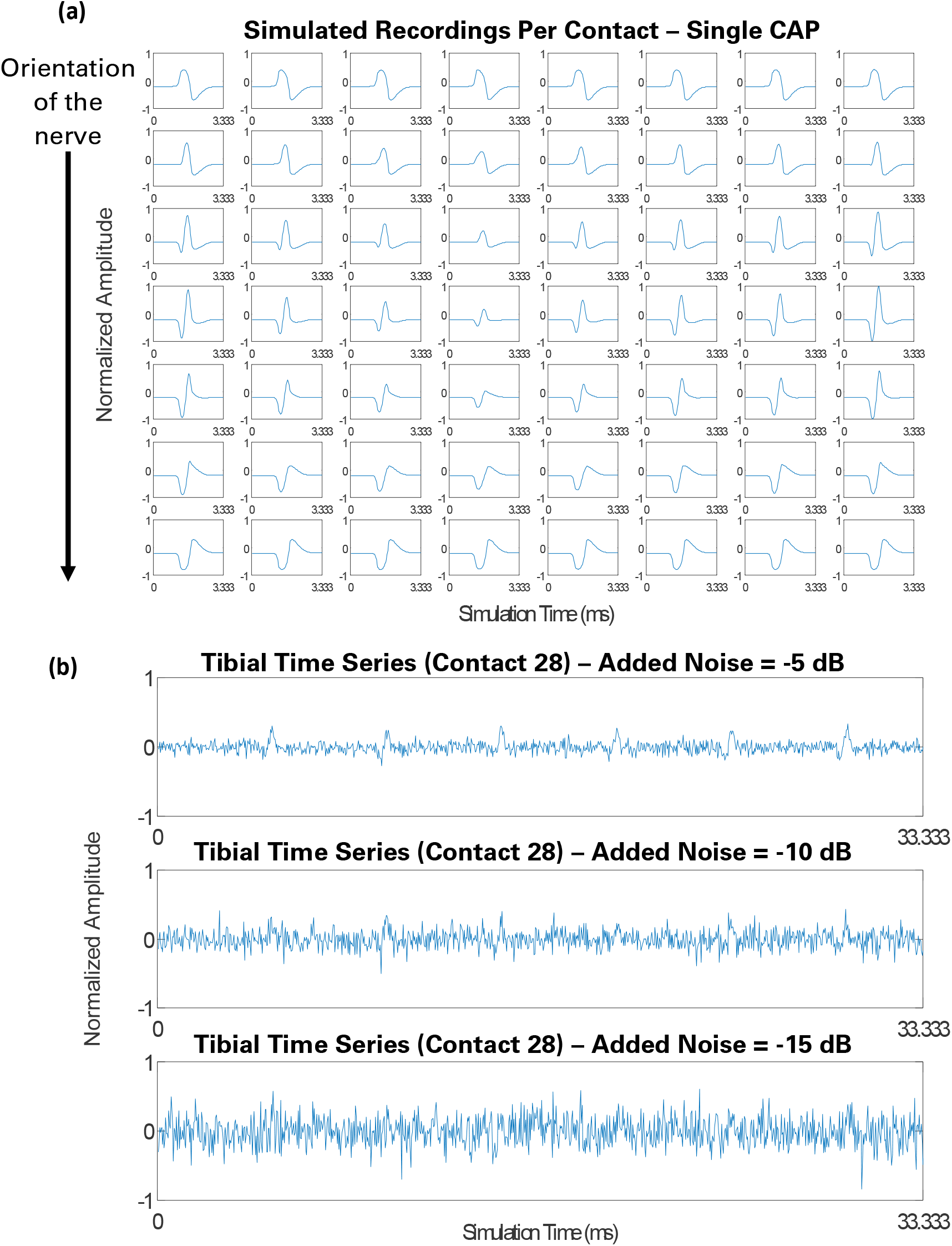
(a) Simulated recordings at all 56 contacts of the cuff electrode for a single CAP propagated through the tibial pathway. (b) Segments of the tibial time series measurements of the middle contact (contact 28) at different noise levels (SNR = −5, −10, and −15 dB). All data here have been normalized for visualization purposes.

### 3.2. Classification

#### 3.2.1. Growth of Encapsulation Tissue

Figure 8 shows the mean F_1_-score obtained by the CNN at each time point in the encapsulation tissue growth simulation using the three updating strategies at decreasing levels of SNR. Note, because of the fact that all folds here were based on very similar synthetic data, the standard deviations were below 0.007, and therefore are not displayed in Figure 8 for ease of visualization.

**Figure 8:**
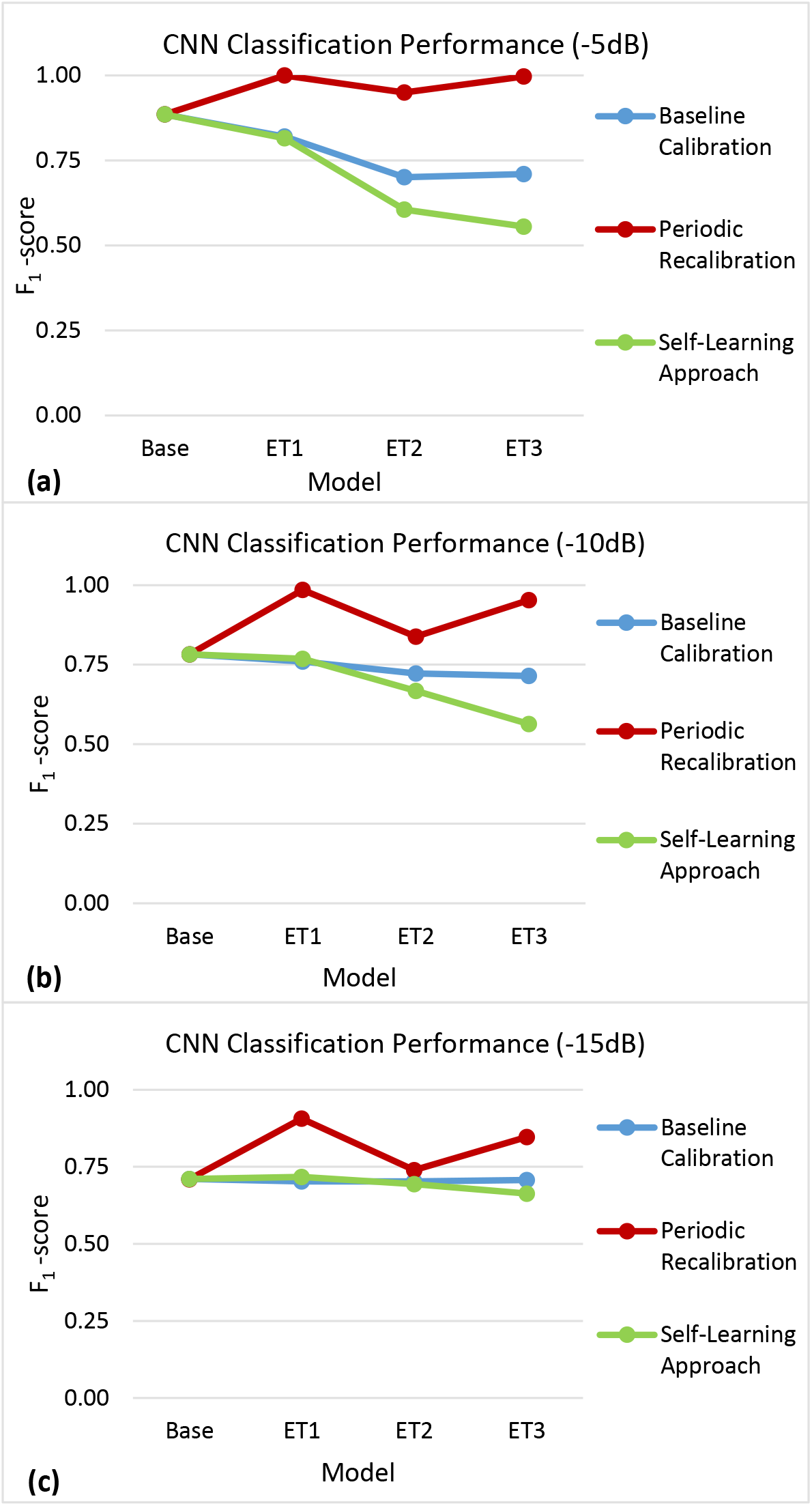
F_1_-scores obtained in the encapsulation tissue simulation using the baseline calibration (blue), periodic recalibration (red), and self-learning (green) update approaches at different time points (represented here as additional models) using measurements with (a) −5 dB, (b) −10 dB, and (c) −15 dB of noise added.

#### 3.2.2. Rotation of the Nerve Cuff Electrode

Figure 9 shows the mean F_1_-score obtained by the CNN at each time point in the rotation of the nerve cuff electrode simulation using the three updating strategies at decreasing levels of SNR. Note, because of the fact that all folds here were again based on very similar synthetic data, the standard deviations were below 0.008, and therefore are not displayed in Figure 9 for ease of visualization.

**Figure 9:**
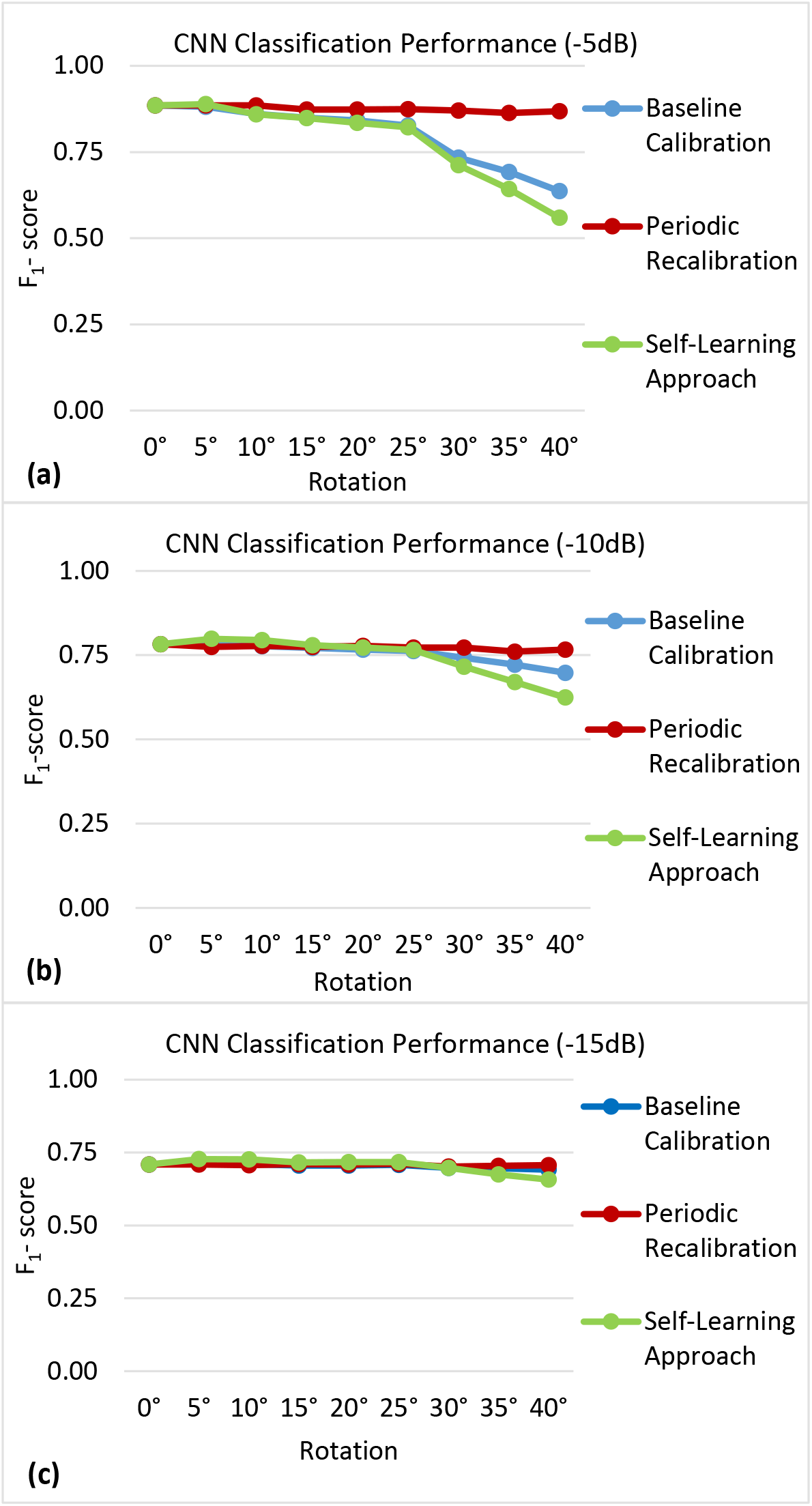
F_1_-scores obtained in the rotation of the cuff electrode simulation using the baseline calibration (blue), periodic recalibration (red), and self-learning (green) update approaches at different time points (represented here as additional models) using measurements with (a) −5 dB, (b) −10 dB, and (c) −15 dB of noise added.

#### 3.2.3. Influence of Training Frequency and Initial Performance on Self-Learning Approach Performance

The classification performance over time of the self-learning approach using different training frequencies and starting at two different initial performance levels for each noise level are shown in Figure 10. In each case, training and evaluation was conducted up until the time point in which there were no longer enough samples (i.e., less than 200) from each class to form a sufficient validation set.

**Figure 10:**
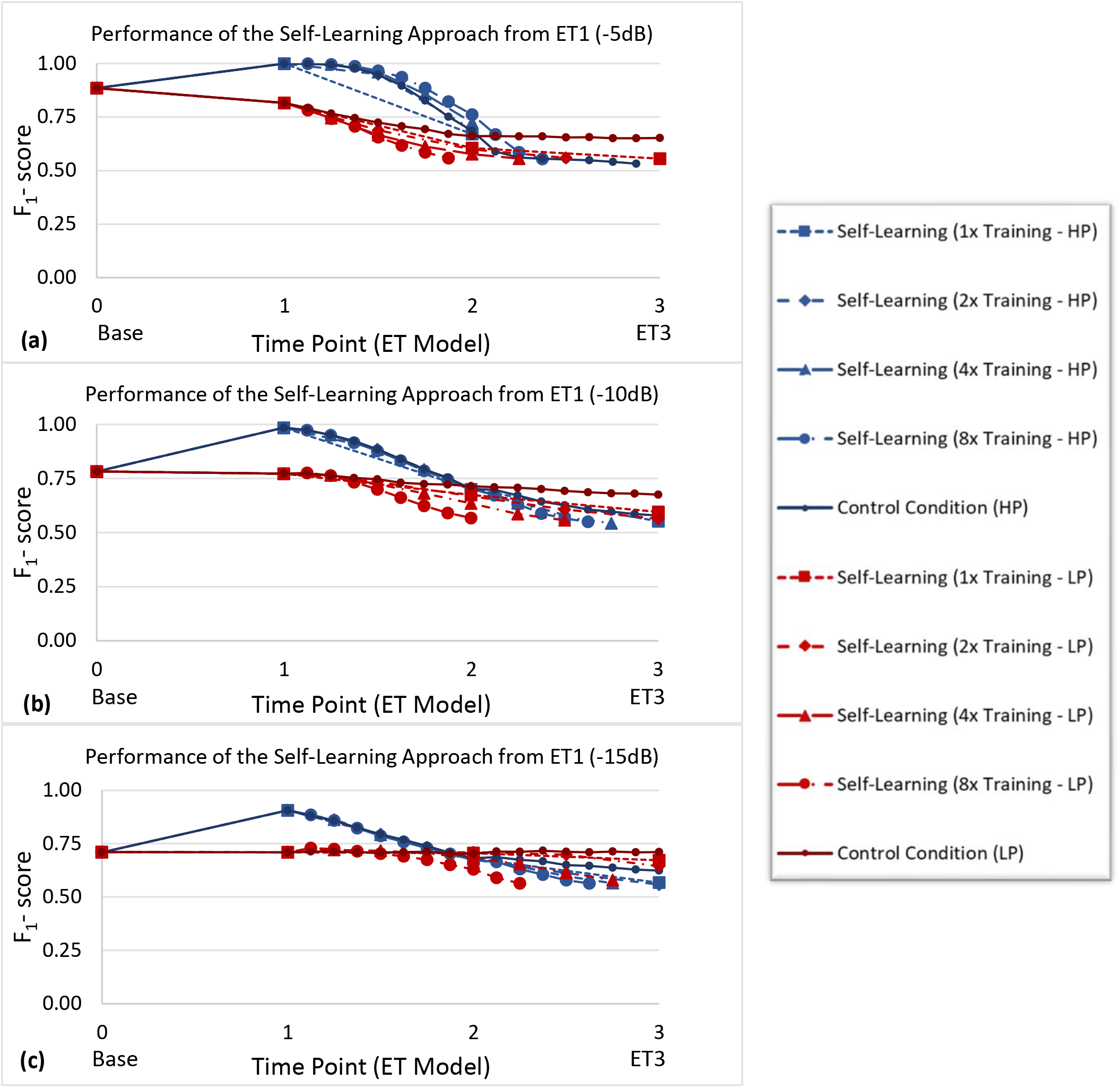
Classification performance of the self-learning approach using different training frequencies and starting at two different initial performance levels for a SNR of (a) −5 dB, (b) −10 dB, and (c) −15 dB. Here, HP represents starting at the Higher Performance (HP) level shown as the blue curves, while LP represents starting at the Lower Performance (LP) level shown as the red curves. The starting point for HP was obtained by applying the periodic recalibration approach between Base and ET1. The starting point for LP was obtained by applying the self-learning approach between Base and ET1.

Figure 11 shows how the rate of decline varies as a function of the training frequency, initial performance level, and noise level. The values represented on these plots were derived by subtracting the slope of the control between ET1 and ET2 from the corresponding slope of the self-learning classifier. Therefore, in Figure 11, red areas represent situations in which the self-learning approach outperformed the control condition, while blue areas represent situations in which the self-learning approach performed worse than the control condition.

**Figure 11:**
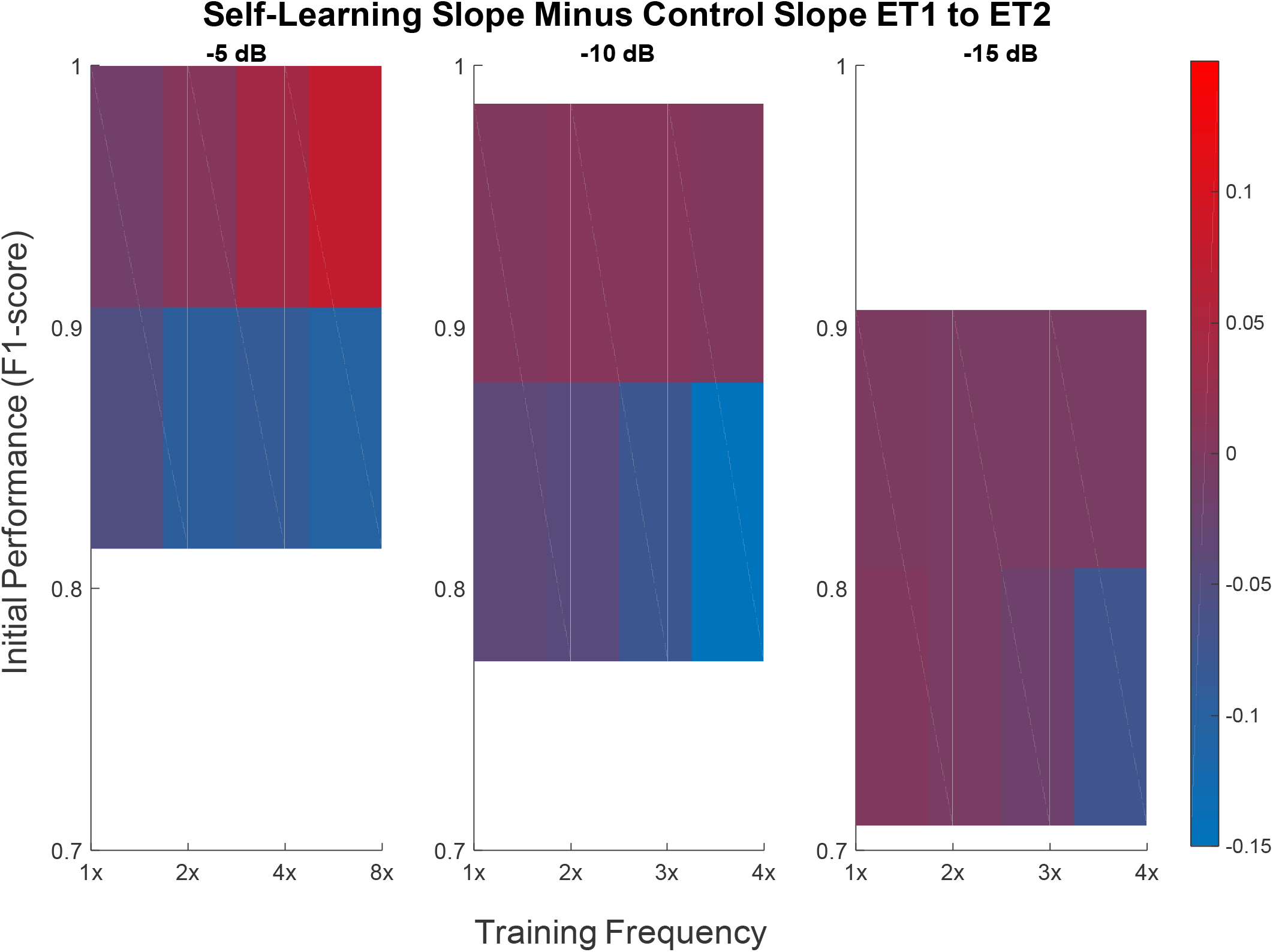
Self-learning slope minus control slope between ET1 and ET2, for a SNRs of −5, −10 and −15 dB. The −5 dB plot (a) shows two levels of initial performance (F_1_-score = 0.9997 and 0.8153). The −10 dB plot (b) shows two levels of initial performance (F_1_-score = 0.9852 and 0.7720). The −15 dB plot (c) shows two levels of initial performance (F_1_-score = 0.9065 and 0.7096).

## 4. Discussion

In this simulation study, we characterized the expected changes in the performance of a selective recording approach during chronic implantations and evaluated strategies to compensate for these changes. We focused on two commonly seen challenges in chronic peripheral nerve interface operation: growth of encapsulation tissue and movement of the nerve cuff electrode. In each of these scenarios, three update strategies were investigated: baseline calibration, periodic recalibration, and a self-learning approach. The baseline calibration approach represented a control condition used to determine the effect of the particular challenge being investigated over time and therefore provided a comparison point for the other two update strategies.

### 4.1. Baseline Calibration

As Figure 8 illustrates, the baseline calibration approach, in which the performance of a static CNN is evaluated at specific time points, begins to degrade as the encapsulation tissue grows. This trend is seen more clearly at higher signal-to-noise ratios of −5 and −10 dB, whereas the performance of the baseline calibration approach is relatively more stable at the lowest SNR of − 15 dB. Nonetheless, a general degradation in classification performance over time is reasonable here, as previous research has shown that the growth of this encapsulating tissue is known to cause changes in the characteristics of the bioelectric signal [43]–[45].

Figure 9 demonstrates relatively similar results, in which a degradation of the selective recording algorithm is most evident when inspecting the baseline calibration at the higher SNRs of −5 and − 10 dB. At these noise levels, the general trend of the network is a decline in classification performance as the rotation of the cuff electrode increases. Again, this result is reasonable, as previous research has shown that a change in the initial position of the nerve cuff electrode will likely cause a modification in the recordings of the bioelectric signals due to the geometry of the volume conductor changing [46], [47], [56].

In summary, in both simulations, we observed that a network initially trained at baseline, which achieves relatively high performance, begins to misclassify new samples as we simulate the gradual growth of encapsulation tissue and progressive electrode movement. In contrast to this observed decrease, the performance of the baseline calibration approach at the lowest SNR of −15 dB was relatively stable in both simulations, albeit at a lower level of performance. It is possible that the noisier training data may have led to a network that is better able to generalize and therefore suffers less of a decline as additional signal distortions occur. Nonetheless, these observations support the hypothesis that selective recording performance will decline over time, calling for strategies to compensate for these changes.

### 4.2. Periodic Recalibration

The periodic recalibration approach showed an improvement in classification performance in the growth of encapsulation tissue simulation (Figure 8) and a maintenance in classification performance in the rotation of the cuff electrode simulation (Figure 9).

An interesting observation is the consistent pattern displayed across all noise levels in the encapsulation tissue simulation. The periodic recalibration performance initially increases at time point ET1, decreases at time point ET2, before finally increasing once again at the final time point, ET3. This result may provide evidence to suggest that the recorded signals from different pathways may be more easily characterized and discriminated when the encapsulation tissue fully fills the space between the epineurium and the cuff electrode, as in time point ET3.

Potential mechanisms for this trend could be the increased CAP amplitude observed in our simulated recordings when the space between the cuff and nerve was filled with encapsulation tissue, and decreased shunting effects from the saline layer. Previously, Moffit et al. modeled a microelectrode array in the presence of encapsulation tissue growth [57] and reported an increase in signal amplitude directly proportional to the thickness of the modeled encapsulation tissue, a result that offered a possible explanation to observations of signal increases seen experimentally by Vetter et al [58]. While referring to a different neural interface, these results are consistent with our finding here and support the validity of our models.

In summary, both simulations provide evidence that a network periodically retrained with new and explicitly labelled samples can adapt effectively to changes in bioelectric signal characteristics during chronic implantation.

Practically, periodic recalibration would require action on the part of the end user, as well as the possible involvement of a clinician or other support personnel, and therefore reduce the convenience of the system. It is difficult to make an accurate estimation of the frequency of recalibration that would be needed for this technique to be effective based on the results of this study, and considering the fact that other chronic studies mentioning nerve cuff electrode encapsulation tissue growth *in vivo* [15], [20], [43] only assessed the degree of encapsulation tissue formation at the end of their studies (anywhere between 9-80 weeks post-implantation).

That being said, however, there is mention of abnormal tissue growth and fiber morphology [20], as well as the modification of the electrical properties associated with nerve cuff electrodes [59] seen as early as two weeks post-implantation. Therefore, it is not unlikely that a recalibration procedure may be required every two to four weeks in order to be effective, at least in the early stages. Considering this fact, employing a periodic recalibration approach would likely lessen the overall usability of a system implementing a neural interface with a selective recording algorithm, which motivates the use of a self-learning approach to minimize human intervention.

### 4.3. Self-Learning Approach

The self-learning approach, in which the CNN is retrained with self-labelled data at specific time points, showed a degradation in classification performance over time, similar to that of the baseline calibration approach, in both simulations. These results can somewhat be expected as the samples used to retrain the network in this approach are labelled by the network itself, and only those with high enough confidence are used to create a training dataset. Because this is the case, the training set used in this approach was often smaller than that of the other two updating approaches. Although the samples incorporated into the self-labelled datasets are based upon confidence level, this does not necessarily prevent the network from including incorrectly labelled samples into the training sets used and as such, an accumulation of incorrectly labelled samples begins to occur over time. This is the most likely explanation for the degradation trend seen in the self-learning approach and indicates that additional refinements to this updating approach are required in order to improve its efficacy. Like the baseline calibration approach, this trend was more obvious at higher signal-to-noise ratios of −5 and −10 dB, whereas the performance of the self-learning approach was relatively more stable at the lowest SNR of −15 dB.

A pattern observed in both the baseline calibration and self-learning approaches is that performance was relatively stable until a rotation of 25°, at which point the performance began to decline more sharply. This trend was consistent across all three noise levels investigated. It is possible that this phenomenon was due to the geometry of the cuff, as contacts are spaced or located at intervals of 45°. Thus, at a rotation of 20°, the current position of each contact is closer to its original location, compared to the original position of another contact. Once rotated 25° or greater, contacts are closer to the original location of some other contact. Because of this, the CNN may misinterpret the spatiotemporal signatures, possibly leading to a decline in performance.

In summary, in both simulations, a network that achieved relatively high performance upon initially training at baseline and then used a self-learning update strategy began to misclassify new samples as we simulated the gradual growth of encapsulation tissue and progressive electrode movement. We therefore conducted additional analysis of the factors that may influence self-learning performance.

### 4.4. Influence of training frequency and initial performance on self-learning approach performance

Two major factors thought to influence the efficacy of the self-learning approach were investigated in this simulation: the frequency of training as well as the initial performance level.

In order to quantify the effects of these factors on this strategy, the self-learning approach was applied at six different initial starting performances across three noise levels. At each of these initial performance levels, each training frequency (one-time, two-times, four-times, and eight-times) was applied, shown in Figure 10.

Figure 10 illustrates a few important points about the self-learning approach. When starting at the higher initial performance level, the difference between the self-learning approach and the control condition decreases along with the SNR. In contrast, for the lower initial performance level, the control condition always outperforms the self-learning approach. Furthermore, in the case of lower initial performance levels, more frequent training generally leads to worse performance over time, supporting the hypothesis of an error accumulation phenomenon.

In the final step of this analysis, Figure 11 shows the relationship between the rate of decline, the initial performance level, the frequency of training, and the level of noise. This data highlights specific cases in which the self-learning approach was able to slow the performance decline over time. Specifically, in the case of recordings of relatively high SNRs in the context of nerve cuff recordings, implementing the self-learning approach at a high initial performance, with frequent retraining, can slow the decline of the selective recording algorithm’s performance over time as a result of encapsulation tissue growth. The implications of this are that a system implementing a self-learning approach may require less frequent retraining, ultimately improving both its convenience and usability.

## 5. Conclusion

This simulation study confirms that the performance of a selective recording approach, based on multi-contact nerve cuff recordings and a CNN trained at baseline, will degrade over time, and demonstrates strategies to compensate for this trend. A periodic recalibration approach can help maintain or even improve selective recording performance over time in the presence of chronic implantation challenges (encapsulation tissue and electrode movement), at the cost of increased demands on the user. A self-learning update approach, in which an algorithm is retrained at specific points in time with new, self-labelled data, will in most case fail over time as a result of an eventual accumulation of mislabeled training data. However, under specific conditions (high SNR, high initial performance, and frequent retraining), this strategy may help to slow the rate of decline of selective recording performance, compared to a control condition. These findings will assist in the development of more intuitive, robust, and longer lasting closed-loop neuroprosthetic and neuromodulation applications. In particular, the practical implications of this research are to form the basis for establishing recalibration schedules, and for implementing automated self-learning techniques that reduce the necessary recalibration frequency.

## 6. Acknowledgements

This work was supported in part by the Canadian Institutes of Health Research, the Toronto Rehabilitation Institute, and the Natural Sciences and Engineering Research Council of Canada (RGPIN-2014-05498, RGPIN-2020-06246).

